# Preoptic area controls sleep-related seizure onset in a genetic epilepsy mouse model

**DOI:** 10.1101/2023.11.24.568593

**Authors:** Cobie Victoria Potesta, Madeleine Sandra Cargile, Andrea Yan, Sarah Xiong, Robert L. Macdonald, Martin J. Gallagher, Chengwen Zhou

**Affiliations:** Department of Neurology, Vanderbilt University Medical Center, Nashville, TN 37232; Speech, Language and Hearing Science Program, Auburn University, Auburn, AL 36849; Vanderbilt Brain Institute and Neuroscience graduate program, Vanderbilt University Medical Center, Nashville, TN 37232

## Abstract

In genetic and refractory epileptic patients, seizure activity exhibits sleep-related modulation/regulation and sleep and seizure are intermingled. In this study, by using one het *Gabrg2^Q390X^* KI mice as a genetic epilepsy model and optogenetic method *in vivo*, we found that subcortical POA neurons were active within epileptic network from the het *Gabrg2^Q390X^*KI mice and the POA activity preceded epileptic (poly)spike-wave discharges(SWD/PSDs) in the het *Gabrg2^Q390X^* KI mice. Meanwhile, as expected, the manipulating of the POA activity relatively altered NREM sleep and wake periods in both wt and the het *Gabrg2^Q390X^* KI mice. Most importantly, the short activation of epileptic cortical neurons alone did not effectively trigger seizure activity in the het *Gabrg2^Q390X^* KI mice. In contrast, compared to the wt mice, combined the POA nucleus activation and short activation of the epileptic cortical neurons effectively triggered or suppressed epileptic activity in the het *Gabrg2^Q390X^* KI mice, indicating that the POA activity can control the brain state to trigger seizure incidence in the het *Gabrg2^Q390X^* KI mice *in vivo.* In addition, the suppression of POA nucleus activity decreased myoclonic jerks in the *Gabrg2^Q390X^*KI mice. Overall, this study discloses an operational mechanism for sleep-dependent seizure incidence in the genetic epilepsy model with the implications for refractory epilepsy. This operational mechanism also underlies myoclonic jerk generation, further with translational implications in seizure treatment for genetic/refractory epileptic patients and with contribution to memory/cognitive deficits in epileptic patients.

## Introduction

Sleep and wake are daily physiological events in animals and humans^1–4^, while sleep spindles in the brain underlie memory consolidation^5–8^. Under pathological conditions in epileptic patients due to such as stroke, brain injury and genetic mutations, sleep can be altered resulting in memory/cognitive deficits^9–16^ in patients. Moreover, sleep disturbance has been associated with genetic and refractory epileptic patients as comorbid symptoms in epileptic patients^9,11,14,15,17,18^. Sleep rhythm seems to modulate epilepsy incidence in refractory epileptic patients^9,11,14,19,20^ and in genetic epileptic animal models^21,22^. Our genetic mouse model with het *Gabrg2^Q390X^*mutation exhibits seizure incident preference during non-rapid eye movement (NREM) sleep^21^. So far, thalamocortical circuitry^23–25^ underlies the circuitry basis to generate sleep spindles and epileptic spike-wave discharges (SWDs) in animal models^10,24,26–28^ and in human pathological conditions^10,29^. However, this thalamocortical circuitry mechanism does not explain epilepsy-caused sleep disturbance or sleep-modulated ictal activity in genetic and refractory epileptic patients, suggesting that other cortical and subcortical circuitry may be involved.

Sleep is controlled by subcortical structures including preoptic area [both extended ventrolateral nucleus (VLPO) and median nucleus (MnPO)^30^, referred as POA together in this manuscript] and ventrolateral periaqueductal gray^1,3,4,31^. These nuclei suppress wake-promoting hypocretin/orexin neurons by their gabaergic neuronal output and cause the brain to switch to NREM sleep state^1,3,31,32^. At the same time, during sleep, cortical neurons within thalamocortical circuitry exhibit up/down-state alteration^7,33–40^. Our previous work indicates that sleep-related up-states (in slow-wave oscillation during NREM sleep) selectively and balanced potentiate excitatory and inhibitory synaptic strength in cortical neurons from wild-type (wt) mice, while sleep-related up-states only potentiate excitatory synaptic strength in cortical neurons from one genetic mouse model with *Gabrg2^Q390X^* mutation^21,41^. Thus, we hypothesize that the subcortical POA can use this synaptic potentiation mechanism to trigger or regulate epilepsy incidence in the het *Gabrg2^Q390X^* knock-in(KI) mice. This mechanism is compatible with one study that stimulating MnPO nucleus can cause absence epilepsy in one rodent epileptic model^42^.

Using one genetic epilepsy mouse model with a *Gabrg2^Q390X^* mutation in gabaergic receptor γ-subunit in cortical neurons and optogenetic method^21,41^, we found that POA multi-unit activity preceded epileptic activity in the het *Gabrg2^Q390X^* KI mice and the POA neurons were active due to epileptic activity in the het *Gabrg2^Q390X^* KI mice. Further, manipulating of the POA activity triggered or suppressed epilepsy incidence in the het *Gabrg2^Q390X^*KI mice, supporting one operational mechanism that the subcortical POA can trigger or regulate epilepsy incidence in this genetic epilepsy model with implications in memory and cognitive deficits in epileptic patients.

## Methods

### Mouse surgery for EEG/optic cannula/tungsten electrode implantation

With all procedures in accordance with guidelines set by the institutional animal care and use committee of Vanderbilt University Medical Center, homozygous cFos-tTA::TetO transgenic mice were created by crossing cFos-tTA mice (#018306 from The Jackson Laboratory)^43^ and TetO mice (#017906, Tg(tetO-hop/EGFP,-COP4/mCherry)6Kftnk/J, from The Jackson Laboratory)^44^. The homozygous mice then crossed with het *Gabrg2^Q390X^* KI mice to breed cFos-tTA::TetO::wt littermates and cFos-tTA::TetO::het *Gabrg2^Q390X^* KI mice (*<u>shortened as wt and het mice unless</u> <u>specified</u>*) to activity-dependently express both halorhodopsin (NpHR) and channelrhodopsin (ChR2) proteins in neurons within cerebral cortex and subcortical structures for the optogenetic manipulation of cortical/subcortical activity. In the cFos-tTA:tetO::wt and cFos-tTA::tetO::het *Gabrg2^Q390X^*KI mice, the NpHR protein expression in the brain was confirmed by checking green GFP expression in neurons and the ChR2 protein expression by red mCherry expression in neurons (further with antibody against mCherry) within brain slices or post-PFA fixed brain sections (see cell counting section). cFos-GFP mice (#014135, from The Jackson Laboratory) were also used to breed cFos-GFP::wt and cFos-GFP::*het Gabrg2^Q390X^*KI mice for tracking cFos-GFP positive neurons within the whole brain. After anesthetized with 1–3% isoflurane (vol/vol) during aseptic surgery, these wt and het mice (both genders, 2-5 month old) were implanted three EEG screw electrodes (each for one hemisphere and one for grounding above cerebellum, Pinnacle Technology, #8201). Meanwhile, depending on experiments, two fiber optic cannulas (0.2-0.4 mm diameter, Thorlabs Inc., Newton, New Jersey) for laser light delivery *in vivo* were also implanted in somatosensory cortex and subcortical preoptic area [POA, extended ventrolateral(VLPO) and median preoptic(MnPO) nuclei^32^] (brain atlas coordinates: somatosensory cortex range, anterior-posterior between –1.82 and –0.46 mm, midline-lateral between +2.0 and +4.0 mm reference to bregma, dorsal-ventral 0.7-1.1 mm reference to pia surface; POA, anterior-posterior between +0.4 and +0.2 mm, midline-lateral between 0.3-0.5 mm reference to bregma, dorsal-ventral 5.5-5.7 mm reference to pia surface). When necessary, EEG screw electrode positions were adjusted for the space of optic cannula implantation. In addition, one bipolar tungsten electrode was implanted into the POA nucleus in some experiments. Tungsten electrode/optic cannula placements within the brain were checked after euthanasia. One pair of EMG leads were inserted into the trapezius muscles to monitor mouse motor activity such as head movements and staring along with simultaneous videos.

### EEG/multi-unit recordings *in vivo*

As the methods before^21^, post-surgery mice were continuously monitored for recovery from anesthesia and remained in the animal care facility for at least 5 days or 1 week (normal wake/sleep circadian rhythm) before simultaneous EEG/multi-unit activity recordings *in vivo,* along with epileptic behavior being video-recorded for Racine-stage scoring. Two-channel EEG (band filtered at 0.1∼100Hz) and one-channel POA multi-unit recordings (band filtered 300-10K Hz) along with one channel EMG(band filtered 10-400Hz) were collected by using two multiClamp 700B amplifiers (total 4 channels and all in current-clamp mode) (Molecular devices Inc., Union City, CA) and Clampex 10 software (Molecular Devices Inc., Union City, CA), and digitized at 20 kHz using a Digidata 1440A. An optic fiber cable was connected to the optic cannula to deliver laser, which was controlled by a yellow DPSS laser for NpHR [MGL-III-589-50 (50mW, Ultralazers Co., Inc)] or a blue DPSS laser for ChR2 [MBL-III-473-100 (100mW, Ultralazers Co., Inc)]. And the laser delivery timing was controlled by Clampex 10 software with continuous 0.5 Hz yellow laser for NpHR activation or 20 Hz blue laser (1 ms for each pulse duration) for ChR2 activation (20-30 ms duration similar as sleep spindle duration^24^). The intensity was determined by slice experiments *ex vivo* (see brain slice section). Single mice could be conducted with both POA activation and suppression with at least 1-2 hour intervals and without specific temporal sequels. Thus, pre-control SWD/PSD data in mice were mixed from fresh start mouse pre-control data and post-optogenetic manipulation mice for summary data.

Mouse epileptic EEG activity was defined as bilateral synchronous (poly)spike-wave discharges(PSDs)(SWDs, 6-12Hz) with duration >1 s and voltage amplitudes at least twofold higher than baseline amplitude. During NREM sleep period, SWD/PSD amplitudes were set as at least 1.5X of baseline amplitude due to large delta EEG amplitude. All atypical and typical absence epilepsy and general tonic-clonic seizures (GTCS) started with SWD/PSDs, accompanied by characteristic motor behaviors. The behavior associated with SWD/PSDs consisted of immobility, facial myoclonus and vibrissal twitching. The SWD/PSDs and animal epileptic behaviors were also checked and confirmed by persons blind to animal genotypes. The onset times of SWD/PSDs were determined by their leading edge points crossing (either upward or downward) the precedent EEG baseline. PSDs will be identified as very brief (< 0.5 second), high frequency (approximately 50 ms interspike interval) complexes^45,46^ with voltages at least twice the background voltage. SWDs have a lower spike frequency (interspike interval 125-166 ms), and a much higher rhythmicity than the PSDs^46^. PSDs often coincide with myoclonical jerks and SWDs. Epileptic myoclonic jerks^46–48^ will be identified as very brief (<1 s), lightening-like body spasms associated with PSDs^49^. Interictal spikes(IS) will be detected by at least 5x standard deviation (with durations <100ms) above baseline EEG mean amplitudes^50,51^ to prevent false positive detection^48^. *Unlike IS, myoclonic seizures(jerks) are associated with head/body/tail-limbs jerks*^48^. Due to the short experiment time of optogenetic manipulation, GTCSs in the het *Gabrg2^Q390X^* KI mice were very rare and not examined in this study. High-frequency tungsten multi-unit activity was obtained by post-experiment band-filtering(400∼800Hz) originally recorded tungsten electrode activity band-filtered at 300-3K Hz *in vivo*. One threshold detection method (at least 2X baseline amplitude) was used to detect high-frequency activity events using Clampfit 10 software (Molecular Devices Inc., Molecular Devices Inc., Union City, CA) and their duration was analyzed. Any high-frequency activity associated with motor behaviors (video monitored) was removed from this analysis.

Sleep period was determined as period without nuchal EMG activity(active sleep) or just twitch activity(quiet sleep), while wake period was defined as large amplitude EMG activity^21^. The spike-sorting method was used to identify multi-unit activity within POA (including both VLPO and MnPO nuclei) during sleep period, by using the spike amplitudes and spike width^52^. These units were confirmed as sleep-selective POA firings and they were further examined whether or not they preceded and/or synchronized with SWD/PSDs. The intervals were defined as the time between the end of the POA multi-unit activity and the adjacent SWD/PSDs (shorten as POA-SWD/PSDs). If multiple SWD/PSDs followed the POA multi-unit activity, we used the first SWD/PSD for the POA-SWD/PSD intervals. In a similar way, if multiple POA multi-unit activities preceded the SWD/PSDs, only the last POA activity were used to determine the intervals. If the POA multi-unit activities synchronized with the SWD/PSDs, we would use the leading point of the POA multi-units and the leading points of the SWD/PSDs.

### cFos-positive puncta/cell counting

Some mice were cardiacally perfused with paraformaldehyde (PFA, 4% in PBS) to fix brain tissues and fixed mouse brains were sectioned in 70µm thickness to examine cFos-GFP positive neurons within somatosensory cortex and subcortex preoptic area regions (which included the implantation area of optic cannulas) from both cFos-GFP::wt (or cFos-tTA:tetO::wt) and cFos-GFP::het *Gabrg2^Q390X^* KI mice (or cFos-tTA::tetO::het *Gabrg2^Q390X^*KI mice). Neurons were also stained with DAPI (VECTASHIELD® mounting media H-1500, Vector Laboratories Inc. Burlingame CA) for nucleus stainings. Under one fluorescence microscope(Leica dm6000B, Leica Microsystems Inc., Buffalo Grove, IL) or Zeiss LSM-710 inverted confocal microscope, the cFos-GFP positive neurons within the same brain sections were be counted within a fixed size counting box. Within mouse brain atlas S1 cortex coordinates between –1.82 and –0.46 mm (anterior-posterior), three coronal sections (for each mouse, anterior-posterior) around the –1.5mm (optic cannula implantation site) were used with 200 mm apart. Within mouse brain atlas POA coordinates between anterior-posterior +0.5∼+0.2 mm that cover the implantation coordinates of optic cannulas, the MnPO nucleus was framed by the top of 3^rd^ ventricle and the ventral side of medium septum^53^ and the VLPO nucleus was framed by the lateral edge of 3^rd^ ventricle and extended 250 µm dorsallly and 300∼800 µm laterally in three coronal sections^32^. These neuron counts (*total within 3 sections/both sides*) were further analyzed with Image-J program (downloaded from NIH website), with different thresholding levels.

### Mouse brain slice preparation and recordings

Brain slice preparation has been described in our previous work^41^. Both wt littermates and het *Gabrg2^Q390X^*KI mice aged P60-150^54^ were used in this study. After mice were deeply anesthetized and decapitated, mouse brains were sectioned with a vibratome(Leica VT 1200S, Leica Biosystems Inc) for coronal brain slices (300-320 µm thickness, containing somatosensory cortex or VLPO/MnPO nuclei) in a sucrose-based ice-cold solution (containing [in mM]): 214 sucrose, 2.5 KCl, 1.25 NaH_2_PO_4_, 0.5 CaCl_2_, 10 MgSO_4_, 24 NaHCO_3_, and 11 D-glucose, pH 7.4) and brain slices were incubated in a chamber at 35-36°C for 40 min with continuously oxygenated ACSF (see below for components). Finally slices remained at room temperature for at least 1 h before electrophysiological recordings at 32°C. By using a Nikon infrared/DIC microscope (Eclipse FN1, Nikon Corp. Inc), whole-cell patch-clamp recordings (voltage-or current-clamp) were made from somatosensory cortical cFos-GFP positive layer V pyramidal neurons or VLPO/MnPO neurons while slices were continuously superfused (flow speed 1-1.5 ml/min) with ACSF (containing (mM): 126 NaCl, 2.5 KCl, 1.25 NaH_2_PO_4_, 2 CaCl_2_, 2 MgCl_2_, 26 NaHCO_3_, 10 D-glucose, pH 7.4) bubbled with 95% O_2_/5% CO_2_. Using an internal solution [consisted of (in mM): 120 K-gluconate, 11 KCl, 1 MgCl_2_, 1 CaCl_2_, 0.6 EGTA, 10 HEPES, 2 Na-ATP, 0.6 Na-GTP, 10 K-creatine-phosphate, pH 7.3 and filled electrodes with resistances of 2∼5 MΩ], sEPSCs were recorded at a holding potential –55.8mV (Cl^-^ reversal potential). The internal solution for sIPSCs contained (in mM): 65 K-gluconate, 65 KCl, 10 NaCl, 5 MgSO_4_, 0.6 EGTA, 10 HEPES, 2 Na-ATP, 0.6 Na-GTP, 10 K-creatine-phosphate, pH 7.3. For action potential recordings, the same internal solution was used as that for sEPSC recordings. Access resistance Ra was continuously monitored and recordings with Ra larger than 25MΩ or 20% change were discarded.

Data were collected by using one multiClamp 700B amplifier (filtered at 2 kHz) and digitized at 20 kHz by using one Digidata 1440A(Molecular Devices Inc., Union City, CA), along with Clampex 10 software (Molecular Devices Inc., Union City, CA). Both sEPSCs and action potentials were analyzed with a threshold detection method (2.5X baseline RMS for sEPSC) by using Clampfit 10.0 software program^55,56^. All detected sEPSC events were checked by eye to ensure that their waveforms had normal rising and decaying phase. All figures were prepared with both Microsoft Excel, SigmaPlot and Adobe Photoshop softwares. Data were expressed as mean ± SEM(standard error). Statistics were performed with student paired t-test or one/two-way ANOVA with p<0.05 as significance.

## Results

### Epilepsy-related cFos-GFP positive neurons are present in somatosensory cortex and preoptic nucleus (POA, both MnPO and VLPO) in het *Gabrg2^Q390X^* KI mice

Sleep has been shown to modulate epilepsy activity in human patients, particularly for refractory epilepsy^9,11,14,19,20^ and genetic epileptic animal models^21,22^. Thus, we used cFos-GFP transgenic mice to examined whether cFos-GFP positive neurons^57,58^ were present within cortex and subcortical structures for epileptic network and sleep circuitry^43,59^. By using cFos::wt and cFos::*Gabrg2^Q390X^* KI mice, we identified the cFos-GFP positive neurons within the somatosensory cortex (epileptic network) and the subcortical POA (sleep circuitry). Compared to wt mice (n=6 mice, cortex S1 wt 20X 438.00±63.29 and extended VLPO wt 20X 278.6±64.35), the het *Gabrg2^Q390X^* KI mice (n=7 mice) exhibited more GFP-positive neurons within the somatosensory cortex (Fig. 1, cortex S1 20X het GFP-positive puncta 684.75±67.81, t-test p=0.021) and the subcortical extended VLPO (Fig. 1, extended VLPO 20X het 518.00±51.10, t-test p=0.016), indicating that sleep-circuitry was more active due to epileptic activity from the het *Gabrg2^Q390X^* mice, compatible with our recent work on the NREM sleep preference of epileptic activity incidence in the genetic *Gabrg2^Q390X^*KI epilepsy model^21^.

**Figure 1.**
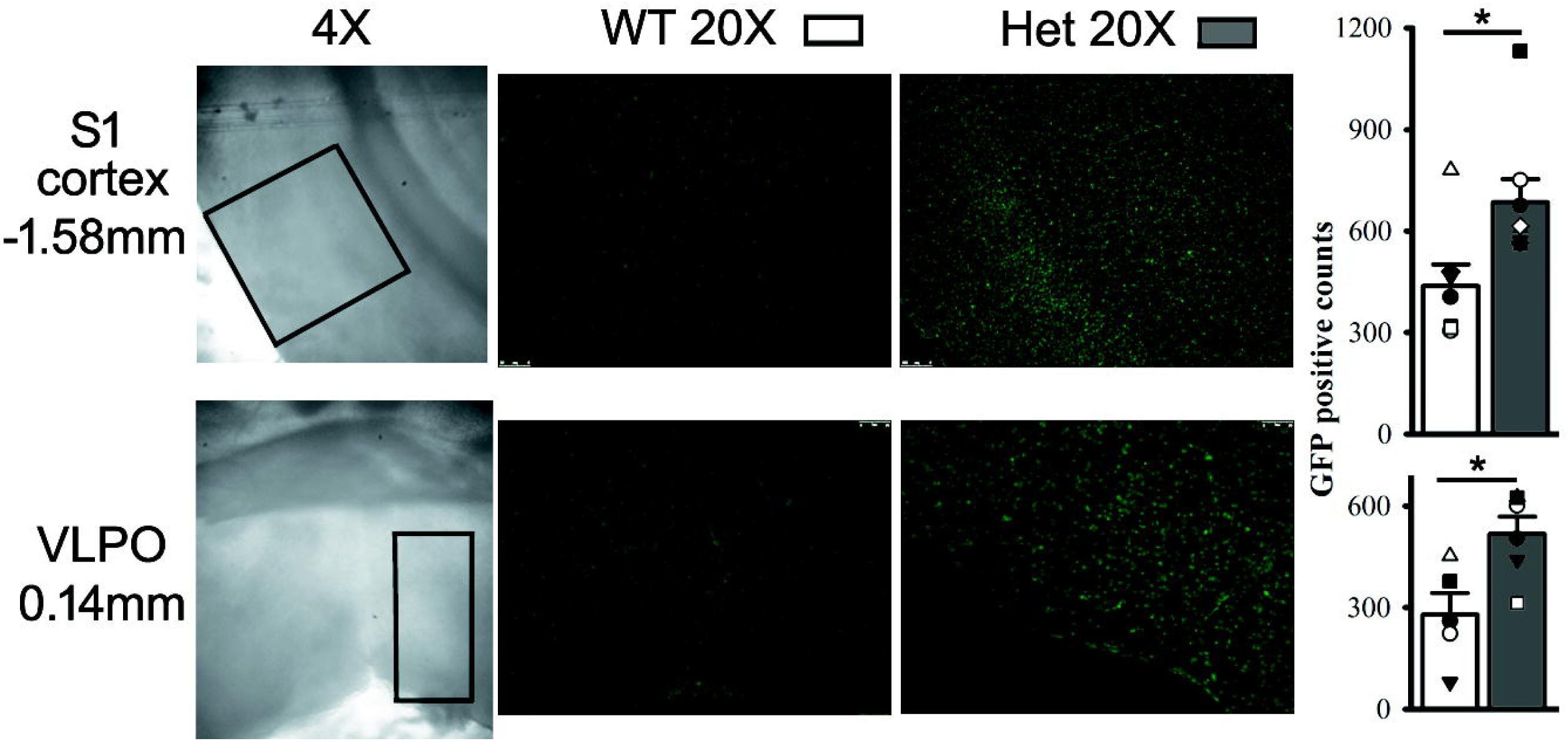
More cFos-GFP positive neurons are present within the preoptic area (POA, MnPO and VLPO) and somatosensory cortex (S1) from het cFos-GFP::*Gabrg2^Q390X^* KI mice. Left panels show 4x DIC images brain sections with counting boxes within the somatosensory cortex (S1 cortex) and the POA and landmarks (hippocampus for S1 cortex and ventral 3^rd^ ventricles/anterior commissure for extended VLPO nucleus^32^), with mouse brain atlas bregma coordinates in the far left. Middle and right panels show wt and het 20x sections (with cFos-GFP positive neurons) containing areas within the counting boxes from the cFos-GFP::wt and het cFos-GFP::*Gabrg2^Q390X^*KI mice (wt n=6 and het n=7 mice) (*counting from total 3 sections/both sides*). * significant change with t-test p<0.05. Otherwise, not significant.

### POA neuronal firings precede epileptic discharges in het *Gabrg2^Q390X^* KI mice

Furthermore, we wanted to examine whether POA neuronal activity was real-time correlated with epilepsy incidence in the het *Gabrg2^Q390X^*KI mice. Multi-unit recordings were used to record neuronal activity within the POA from mice and single-unit activity was spike-sorted/isolated^53^ for those neurons being active during NREM/REM sleep and even wake period. In both the wt and het *Gabrg2^Q390X^*KI mice, some POA neurons were selectively active during NREM sleep (wt 2.19±0.37 Hz, n=8 from 4 mice; het 3.00±0.509 Hz, n=9 from 5 mice) and much less active during REM sleep (wt 0.06±0.04 Hz; het 0.02±0.02 Hz) and wake period (wt 0.50±0.14 Hz; 0.32±0.08 Hz) (Fig. 2A/C), suggesting that these POA neurons were sleep-related and involved in sleep regulation^32,53,60^. In addition, some POA neuronal firings do not show sleep or wake preference (wt n=3, het n=4).

**Figure 2.**
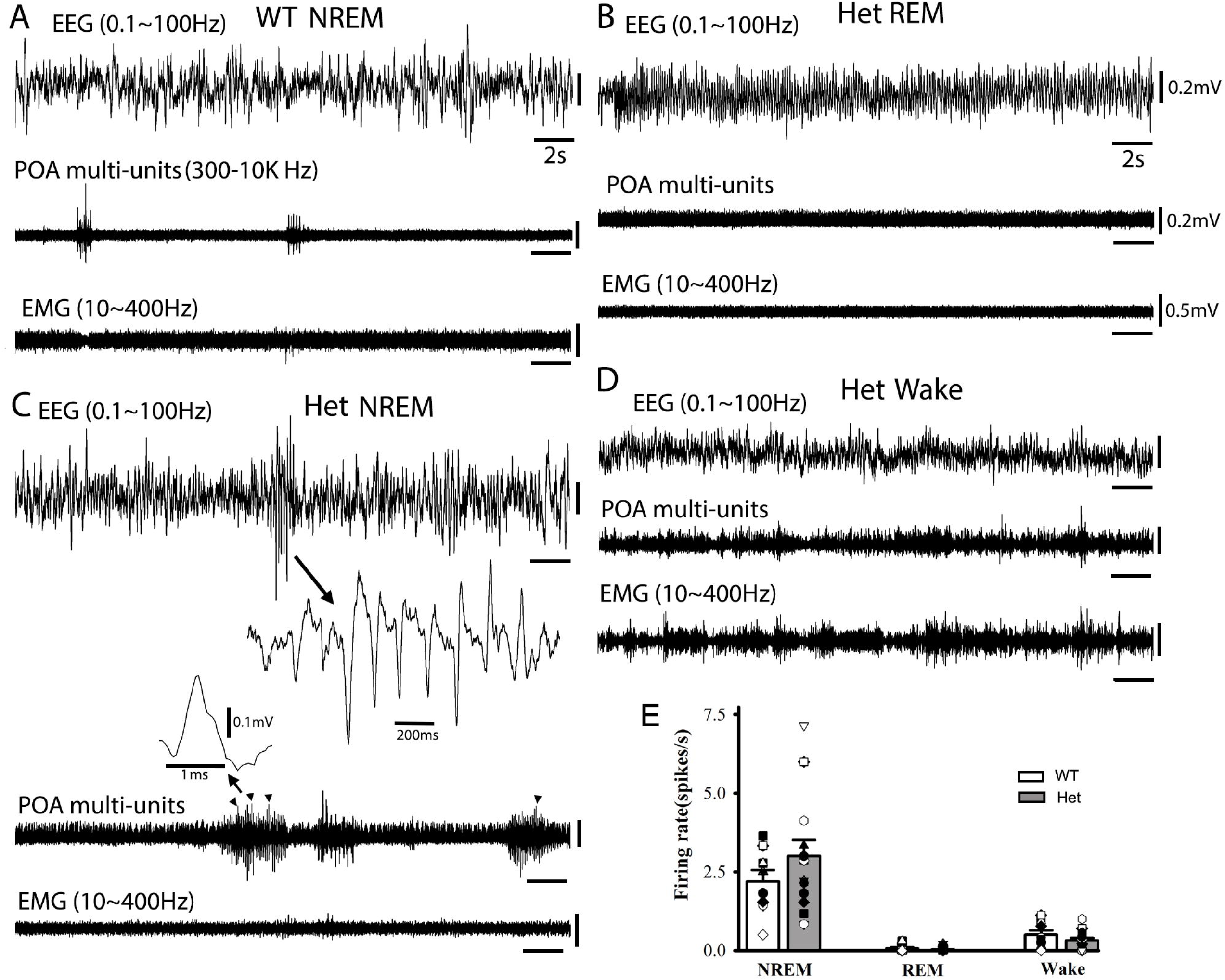
POA neuron firings precede epileptic SWD/PSDs in het *Gabrg2^Q390X^* KI mice during NREM sleep. Panels A-D show simultaneous representative EEG (band filtered 0.1-100Hz)/POA multi-unit tungsten electrode recordings (band filtered 300-10KHz) and EMG recordings (band filtered 10-400Hz) from wt and the het *Gabrg2^Q390X^*KI mice during NREM sleep, REM sleep and wake periods. In panel C, one SWD event is expanded in small temporal scale form EEG traces and one isolated single-unit before/during the SWD event is expanded in small temporal/amplitude scales for spike-sorting method. The arrowheads in the POA multi-unit trace indicate the same detected single-unit firings from the het *Gabrg2^Q390X^* KI mice during NREM sleep. Scale bars are indicated as labeled. Panel E shows firing change of the same spike-sorted single-units across NREM/REM sleep and wake periods (n=5 single-units each from wt and het mice)(n=4, wt and n=5 het mice). * significant change with one-way ANOVA.

In the het *Gabrg2^Q390X^*KI mice, some POA neurons were active and immediately preceded (627.09±150.60 ms, n=7 from 5 het mice) or synchronized with ictal SWD/PSDs discharges (Fig. 2C arrowheads), indicating that these POA neurons were likely involved in the ictal discharge incidence in the het *Gabrg2^Q390X^* KI mice.

### Sleep and arousal/awake states can be altered by sole optogenetic manipulation of POA nucleus *in vivo* in mice

To establish the causal role of the precedent POA neurons in sleep state and further in epilepsy incidence from het *Gabrg2^Q390X^* KI mice, we crossed the het *Gabrg2^Q390X^*KI mice with cFos-tTA::TetO mice^43^ to express ChR2 or NpHR proteins in cFos-GFP/mCherry positive neurons within the cortex and subcortical POA from the wt and het *Gabrg2^Q390X^* KI mice. The cFos-GFP protein expression patterns in the het cFos-tTA::TetO::*Gabrg2^Q390X^* KI mice were very similar to the those in the het cFos::*Gabrg2^Q390X^* mice (Fig. 3A het). By delivering blue laser light, cFos-GFP/mCherry positive neurons within brain slices exhibited action potential firings riding upon depolarization ramp (n=5 neurons from the wt or het tTA::TetO::*Gabrg2^Q390X^*KI mice)(Fig. 3B), while by delivering yellow laser light, the cFos-GFP/mCherry positive neurons within brain slices exhibited no action potential firings upon depolarization ramp (n=6 neurons from the wt or het cFos-tTA::TetO::*Gabrg2^Q390X^* KI mice)(Fig. 3C), confirming that blue or yellow laser lights could effectively activate or suppress cortical/subcortical neurons in the brain from the mice.

**Figure 3.**
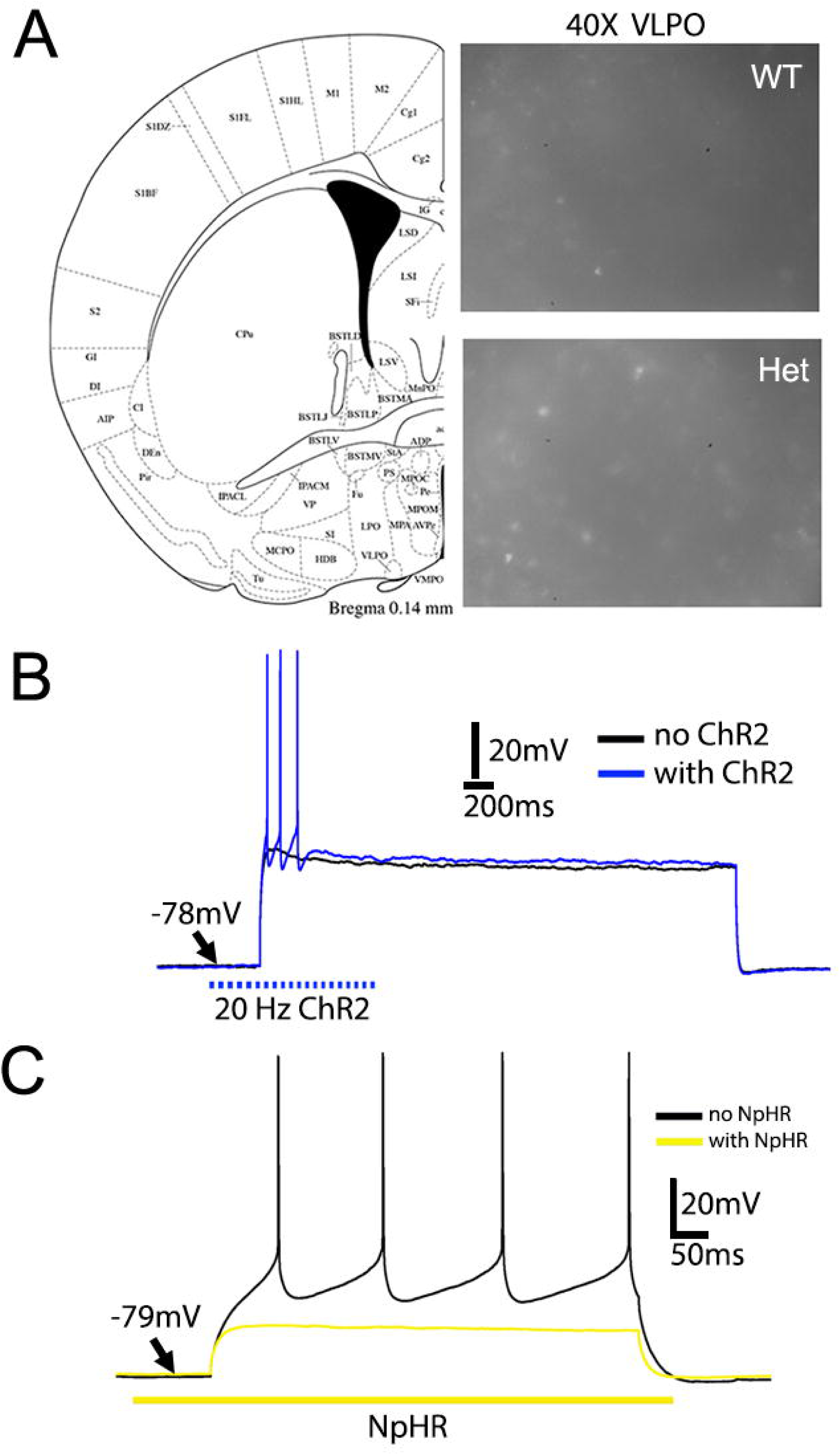
cFos-GFP positive neurons in POA (VLPO) neurons and somatosensory cortical neurons from cFos-tTA::tetO::wt and het cFos-tTA::tetO::*Gabrg2^Q390X^* KI mice. Panel A shows the brain atlas for VLPO (left) and cFos-GFP positive neurons (40X GFP) within VLPO nucleus from wt and the het *Gabrg2^Q390X^* KI mice with more GFP-positive neurons in the het mice. Panel B shows that ChR2-activation by 20 Hz Blue laser (473nm, 1 ms duration) evokes action potentials riding on the depolarization ramp (n=5 cortical neurons from wt or het mice). Panel C shows that NpHR-activation by continuous yellow laser (590nm) completely suppresses the action potential generation by depolarization pulse (n=6 cortical neurons from wt and het mice). Scale bars are indicated as labeled.

Furthermore, with optic cannulas implanted within the POA from the cFos-tTA::TetO::wt and het cFos-tTA::TetO::*Gabrg2^Q390X^*KI mice, we examined whether activation or suppression of the POA could causally alter brain states in the wt or het *Gabrg2^Q390X^* KI mice. As we expected, in both the wt and het *Gabrg2^Q390X^*mice, the blue-laser activation of the POA by expressed ChR2 channels (POA-ChR2), the mice significantly exhibited more NREM sleep/less wake period (Fig. 4A/C) [wt (n=7 mice) NREM pre 62.42±5.45% *vs* post 73.53±4.99%, paired t-test p=0.022; REM pre 0.28±0.23% *vs* post 1.39±0.89%, paired t-test p=0.24, wake pre 37.30±5.49% *vs* post 25.08±5.25%, paired t-test p=0.018; het (n=10 mice) NREM pre 79.12±1.47% *vs* post 83.56±1.48%, paired t-test p=0.036; REM pre 3.5±1.02% *vs* post 5.83±2.23%, paired t-test p=0.306; wake pre 17.38±2.33% *vs* post 12.00±1.69%, paired t-test p=0.042]. In contrast, the yellow-laser suppression of the POA by expressed NpHR channels (POA-NpHR), the mice significantly exhibited less NREM or/and more wake period (Fig. 4B/D) [wt (n=7 mice) NREM pre 74.13±3.38% *vs* post 64.65±4.29%, paired t-test p=0.0006; REM pre 3.23±1.55% *vs* post 6.18±1.66%, paired t-test p=0.027, wake pre 22.64±3.98% *vs* post 29.17±5.10%, paired t-test p=0.012; het (n=10 mice) NREM pre 83.15±1.87% *vs* post 78.57±2.14%, paired t-test p=0.043; REM pre 4.40±1.41% *vs* post 5.33±0.78%, paired t-test p=0.60; wake pre 11.74±1.59% *vs* post 14.58±2.50%, paired t-test p=0.394]. Only REM sleep in the wt mice seemed to be increased by POA-NpHR activation, which might be due to our recordings during daytime period (mouse sleeping period).

**Figure 4.**
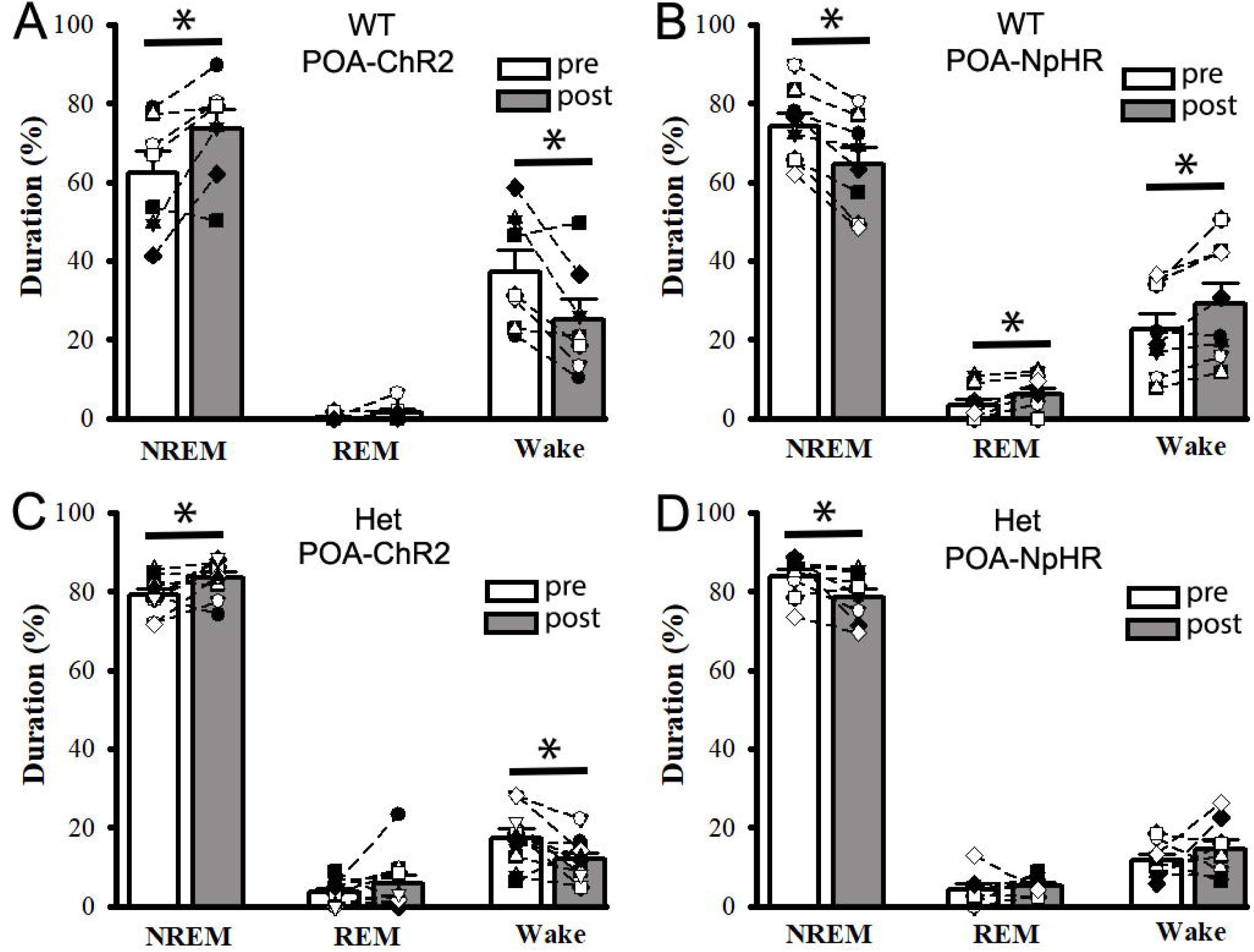
Optogenetic activation/suppression of POA nucleus alone alters NREM/REM sleep and wake periods in wt and het *Gabrg2^Q390X^* KI mice. Panels A/B show the relative changes of NREM/REM sleep and wake period following POA-ChR2 activation (n=8 mice) or POA-NpHR suppression in the wt mice (n=7). Panels C/D show the relative changes of NREM/REM sleep and wake periods following POA-ChR2 activation (n=10 mice) or POA-NpHR suppression in the het *Gabrg2^Q390X^* KI mice (n=8). * significant change with paired t-test p<0.05. Otherwise, paired t-tests are not significant.

### Sole short optogenetic activation of somatosensory cortex does not effectively trigger epileptic activity *in vivo* in het *Gabrg2^Q390X^* KI mice

Our previous work has indicated that sleep-related up-states (in slow-wave oscillations) induced synaptic potentiation in cortical neurons can trigger epileptic activity in het *Gabrg2^Q390X^*KI mice *in vivo*^41^. Thus, we examined whether the manipulation of subcortical POA activity could interact with cortical epileptic neurons within epileptic network to generate ictal activity in the het *Gabrg2^Q390X^*KI mice. By using cFos-tTA::TetO::wt and het cFos-tTA::TetO::*Gabrg2^Q390X^* KI mice, we activated epileptic cortical cFos-GFP/mCherry neurons within somatosensory cortex (shortened as S1-ChR2) from mice in a very short period (30 ms, simulating sleep spindles duration in cortical neurons^24,41^) through blue-laser delivery. In the wt mice, we did not find any epileptic activity, while we did observe epileptic activity in the het *Gabrg2^Q390X^* KI mice. However, blue laser delivery did not significantly evoke epileptic activity in the het *Gabrg2^Q390X^* KI mice [supplemental Fig. 1, wt (n=7 mice) S1-ChR2: SWD/PSDs/hr pre 7.14±3.71 *vs* post 5.57±2.69, paired t-test p=0.18; SWD/PSD duration seconds(s) pre 11.17±5.84 *vs* post 9.52±4.99, paired t-test p=0.19; het (n=7 mice) S1-ChR2: SWD/PSDs/hr pre 66.14±16.07 *vs* post 85.14±23.06, paired t-test p=0.08; SWD/PSD duration(s) pre 228.49±87.89s *vs* post 307.12±114.78, paired t-test p=0.063]. This suggests that without the subcortical POA activity, sleep-spindle like short activation of epileptic cortical neurons within somatosensory cortex is not sufficiently to trigger epileptic activity in the het *Gabrg2^Q390X^*KI mice.

### Combined optogenetic manipulation of the subcortical POA activity and somatosensory cortex activation regulates epilepsy incidence *in vivo* in het *Gabrg2^Q390X^* KI mice

However, when the manipulation of the POA activity was combined with short cortical activation by S1-ChR2 in the het *Gabrg2^Q390X^*KI mice, epileptic activity in the het *Gabrg2^Q390X^* KI mice could be significantly influenced, while combined POA ChR2-activation/NpHR-suppression and cortical S1-ChR2 activation in the wt mice did not show any effect on epileptic activity except sleep or wake period alteration, similar to results as Fig. 4 (in wt)[Fig. 5, wt (n=8 mice) POA-ChR2/S1-ChR2: SWD/PSDs/hr pre 8.625±3.34 *vs* post 7.75±3.13, paired t-test p=0.317; SWD/PSD duration(s) pre 13.28±5.25 *vs* post 11.28±4.78, paired t-test p=0.053] [Fig. 6, wt (n=8 mice) POA-NpHR/S1-ChR2: SWD/PSDs/hr pre 12.5±5.38 *vs* post 12.91±5.43, paired t-test p=0.791; SWD/PSD duration(s) pre 25.41±11.34 *vs* post 20.02±9.49, paired t-test p=0.379]. Specifically, from the het *Gabrg2^Q390X^*KI mice, when the POA was ChR2-activated along with a short cortical S1-ChR2-activation, epileptic SWD/PSD incidence was significantly increased with longer duration, compared to prior epileptic events before laser delivery [Fig. 5, het *Gabrg2^Q390X^*KI mice (n=11 mice) POA-ChR2/S1-ChR2: SWD/PSDs/hr pre 78.38±14.19 *vs* post 132.27±22.60, paired t-test p=0.001; SWD/PSD duration(s) pre 221.56±54.25 *vs* post 538.43±148.09, paired t-test p=0.026]. Furthermore, from het *Gabrg2^Q390X^* KI mice, when the POA was NpHR-suppressed along with the short cortical S1-ChR2-activation, epileptic SWD/PSD incidence was significantly reduced with shorter duration, compared to prior epileptic events before laser delivery [Fig. 6 het *Gabrg2^Q390X^*KI mice (n=9) POA-NpHR/S1-ChR2: SWD/PSDs/hr pre 129.22±34.61 *vs* post 74.56±14.98, paired t-test p=0.029; SWD/PSD duration(s) pre 490.96±157.26 *vs* post 249.89±60.60, paired t-test p=0.043]. Moreover, the effect of epileptic incidence caused by the combined activation of the POA and the cortex S1-ChR2 in the het *Gabrg2^Q390X^* KI mice could be reversed by combined the POA-suppression with cortical S1-ChR2 or in reverse, compared to the precedent POA-activity (see Fig.5G/6G or 5H/6H), indicating that the POA activity can dynamically controls the cortical epileptic activity in the het *Gabrg2^Q390X^*KI mice, compatible with our previous findings and other study.

**Figure 5.**
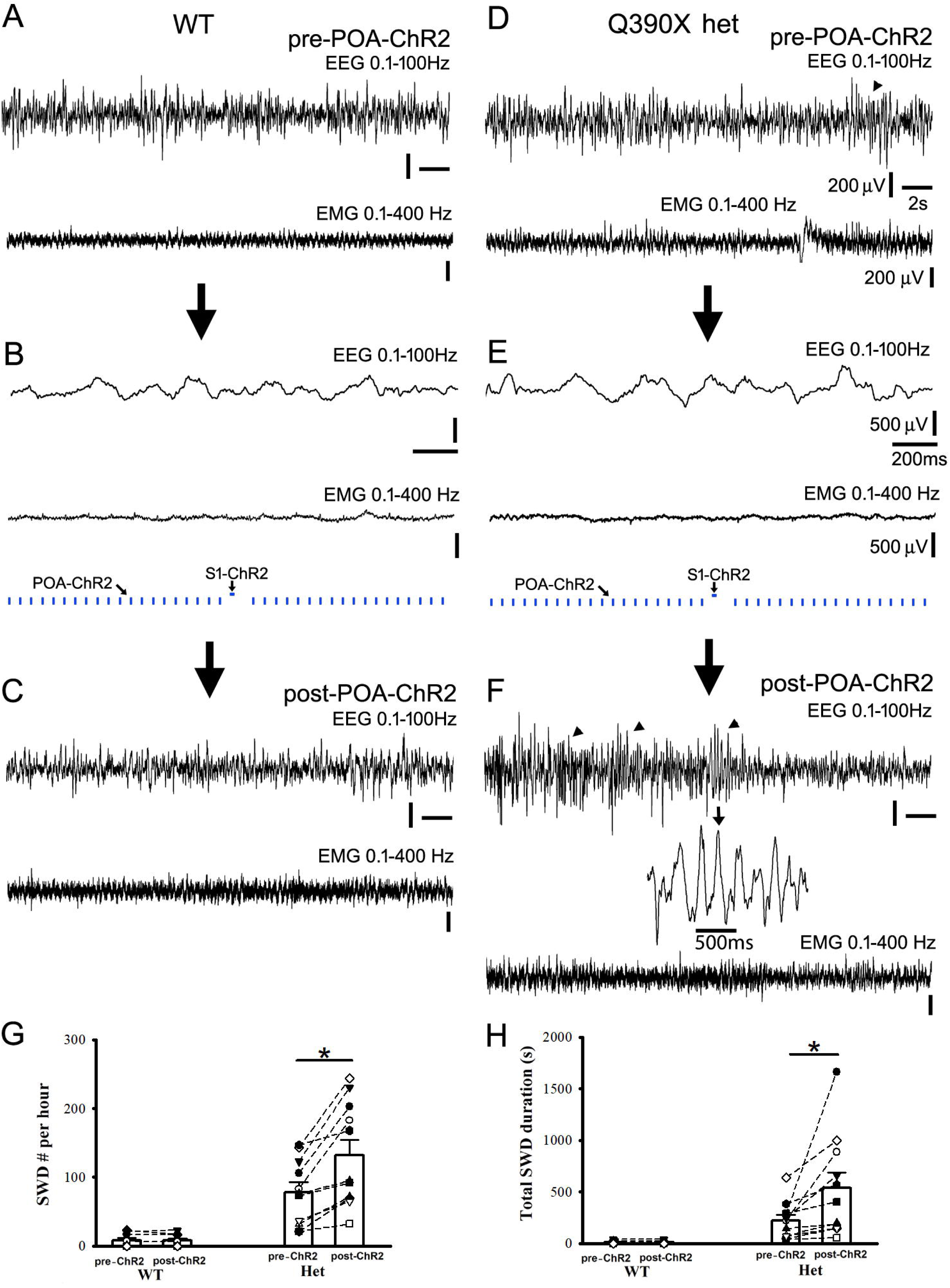
ChR2-activation of POA nucleus along with short activation of cortical neurons increases epileptic SWD/PSDs and duration in het *Gabrg2^Q390X^* KI mice. Panels A/C and D/F show simultaneous representative EEG (band filtered 0.1-100Hz)and EMG recordings (band filtered 10-400Hz) from the wt and the het *Gabrg2^Q390X^* KI mice. Panels B/E show the combined POA-ChR2 activation and somatosensory cortex S1-ChR2 short activation for down/up-states within slow-wave oscillation in the brain from the wt and het *Gabrg2^Q390X^* KI mice *in vivo*. In panel F, one SWD event is expanded in small temporal scale from EEG traces. The arrows between panels show experimental procedure and arrowheads in EEG traces (panels D/F) indicate detected epileptic SWD/PSDs from the het *Gabrg2^Q390X^* KI mice. Scale bars are indicated as labeled. Panel G/H shows summary data of epileptic SWD/PSDs and their duration changes following POA-ChR2/S1-ChR2. In panel G, one inset shows the experimental design for laser delivery. Wt n=8 mice and het n=11 mice. * significant change with paired t-test p<0.05. Otherwise, paired t-tests are not significant.

**Figure 6.**
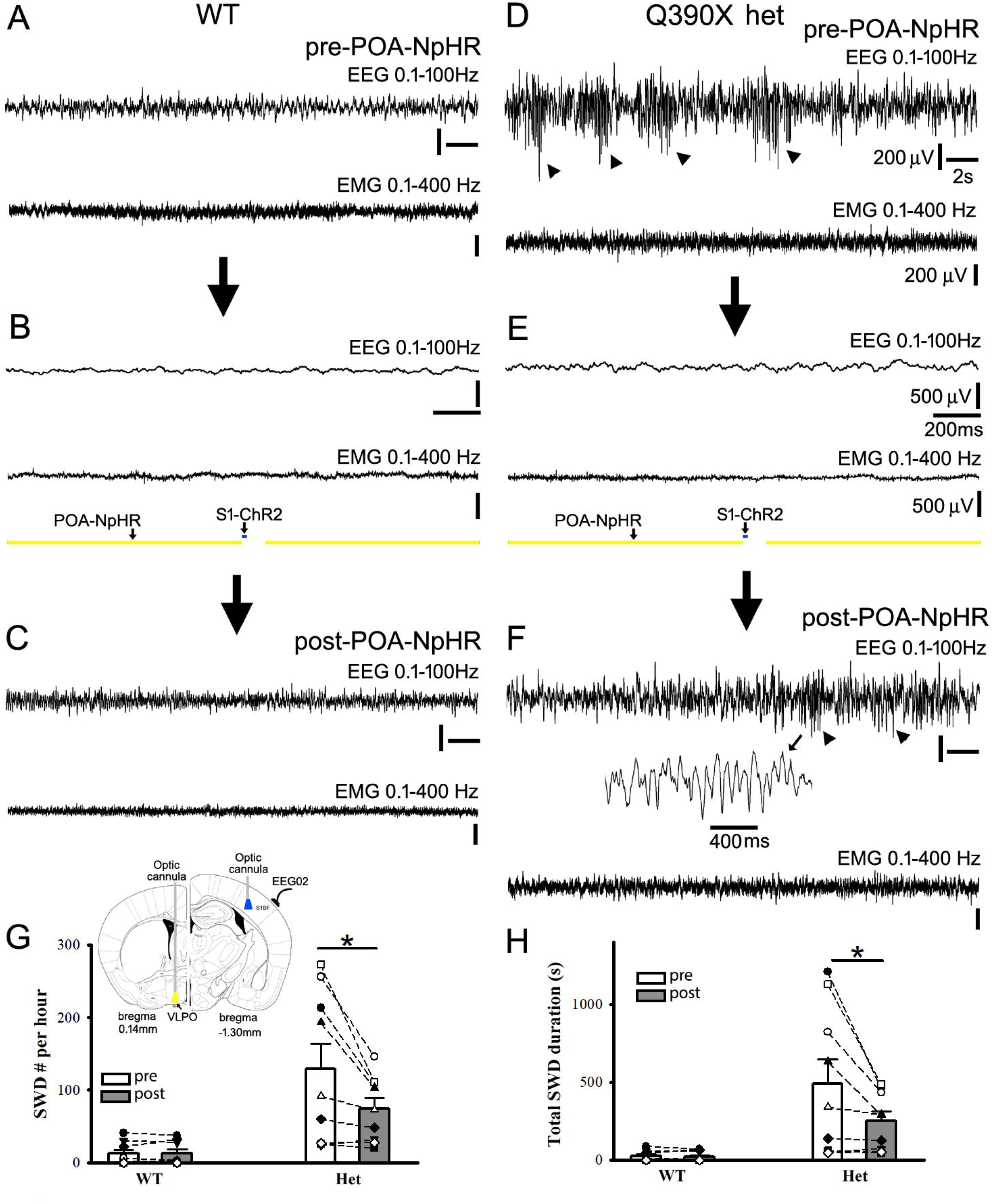
Suppression of POA nucleus by NpHR along with short activation of cortical neurons decreases epileptic SWD/PSDs and duration in het *Gabrg2^Q390X^* KI mice. Panels A/C and D/F show simultaneous representative EEG (band filtered 0.1-100Hz) and EMG recordings (band filtered 10-400Hz) from the wt and the het *Gabrg2^Q390X^* KI mice. Panels B/E show the combined POA-NpHR suppression and S1-ChR2 activation in the brain from the wt and het *Gabrg2^Q390X^*KI mice *in vivo*. In panel F, one SWD event is expanded in small temporal scale from EEG traces. The arrows between panels show experimental procedure and arrowheads in EEG traces (panels D/F) indicate detected epileptic SWD/PSDs from the het *Gabrg2^Q390X^* KI mice. Scale bars are indicated as labeled. Panel G/H shows summary data of epileptic SWD/PSDs and their duration changes following POA-NpHR/S1-ChR2 manipulation. In panel G, one inset shows the experimental design for laser delivery. Wt n=8 mice and het n=9 mice). * significant change with paired t-test p<0.05. Otherwise, paired t-tests are not significant.

In addition, sleep spindle generation depends on brain sleep state and cortical neuron ensemble activation. With combined POA-ChR2 and S1-ChR2 optogenetic manipulation in wt littermate mice *in vivo*, sleep spindle density was increased (supplemental Fig. 1 wt, pre 0.0095±0.0010 vs post 0.0170±0.0020, n=6 mice, paired t-test p=0.015), while in het *Gabrg2^Q390X^* KI mice, sleep spindle density was reduced (supplemental Fig. 1 het, pre 0.0081±0.0007 vs post 0.0060±0.0005, n=11 mice, paired t-test p=0.015), indicating that the generation of epileptic activity may occlude the generation of sleep spindles in in het *Gabrg2^Q390X^* KI mice.

Meanwhile, we did not observe the effect of POA activation (except suppression) on myoclonic jerk activity(MJs)^46^ in the het *Gabrg2^Q390X^*KI mice, suggesting that myoclonic jerks may have different incident mechanism^47,48^ than SWD/PSDs in the het *Gabrg2^Q390X^*KI mice[Fig. 7, wt, (n=8 mice) POA-ChR2/S1-ChR2: MJs/hr pre 2.37±0.94 *vs* post 3.75±1.47, paired t-test p=0.073; POA-NpHR/S1-ChR2: MJs/hr pre 2.75±1.11 *vs* post 2.88±1.20, paired t-test p=0.59] [het *Gabrg2^Q390X^*KI mice (n=9 mice) POA-ChR2/S1-ChR2: MJs/hr pre 11.25±2.55 *vs* post 11.75±1.89, paired t-test p=0.789; (n=10 mice) POA-NpHR/S1-ChR2: MJs/hr pre 13.56±3.27 *vs* post 9.44±2.02, paired t-test p=0.042].

**Figure 7.**
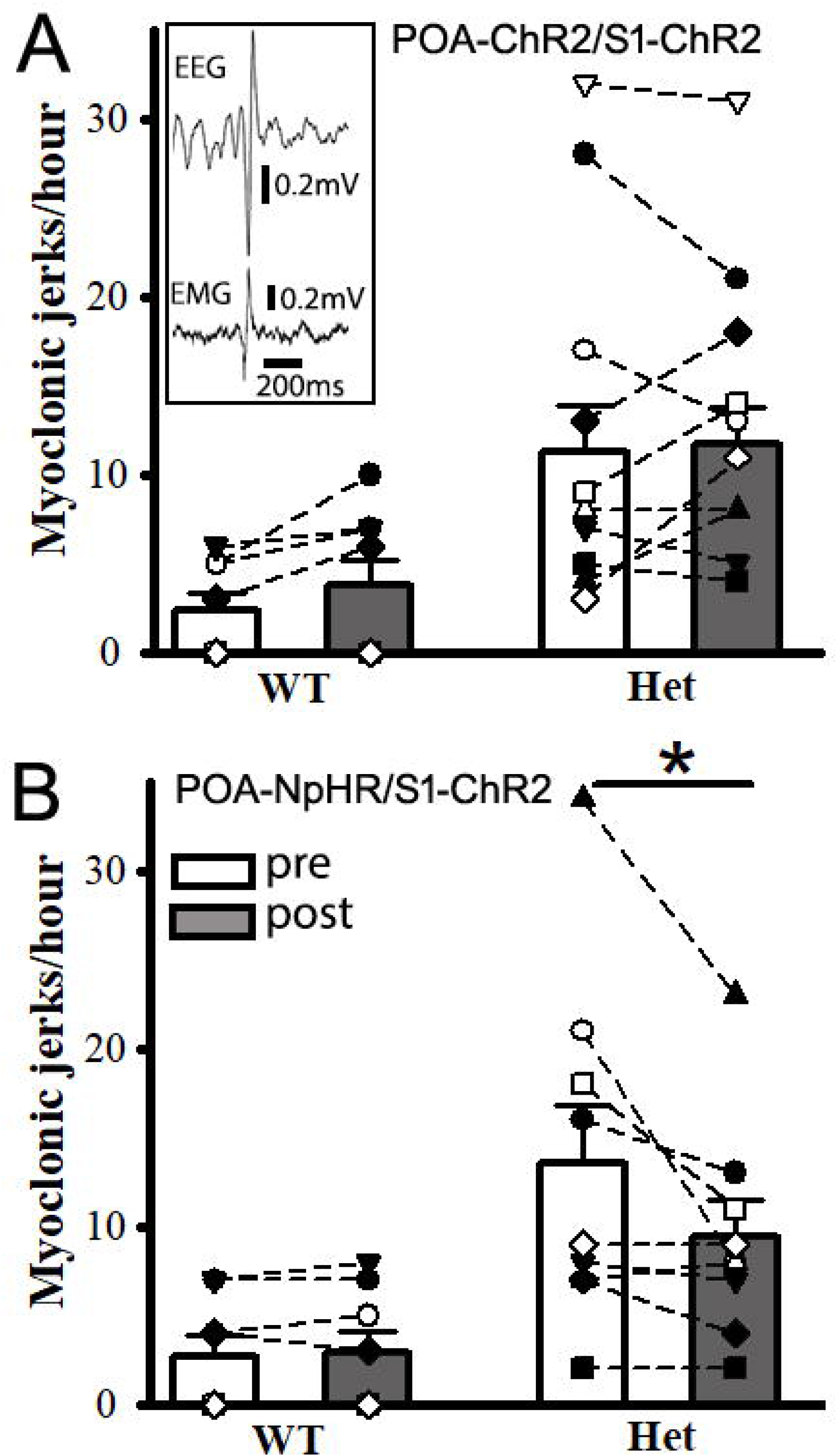
Suppression of POA nucleus along with short activation of cortical neurons decreases myoclonic jerks in het *Gabrg2^Q390X^* KI mice. Panel A shows summary myoclonic jerk changes following POA-ChR2/S1-ChR2 activation. The inset shows the one representative myoclonic jerk with EEG/EMG traces. Scale bars are indicated as labeled. Wt n=8 mice, het n=10 mice. No significant change with paired t-test. Panel B shows summary myoclonic jerk changes following POA-NpHR suppression and S1-ChR2 activation. Wt n=8 mice, het n=9 mice. * significant change with paired t-test p<0.05. Otherwise, paired t-tests are not significant.

## Discussion

In this study we have found that the subcortical POA neurons were active within epileptic network from the het *Gabrg2^Q390X^* KI mice and the POA activity preceded the epileptic SWD/PSDs in the het *Gabrg2^Q390X^* KI mice. Meanwhile, manipulating of the POA activity could alter NREM sleep and wake periods in both wt and het *Gabrg2^Q390X^* KI mice. Most importantly, the short activation of epileptic cortical neurons alone did not effectively to trigger seizure activity in the het *Gabrg2^Q390X^* KI mice. In contrast, combined the POA nucleus activation or suppression along with the short activation of epileptic cortical neurons effectively enhanced or suppressed epileptic activity in the het *Gabrg2^Q390X^* KI mice, indicating that the POA activity can alter the brain state to effective trigger seizure activity in the het *Gabrg2^Q390X^* KI mice *in vivo,* plus to influence myoclonic jerk activity in the het KI mice. Overall, this study discloses an operational mechanism for sleep-related seizure incidence in this genetic epilepsy model, also for refractory epilepsy.

In this study, the data support an operational mechanism by which subcortical POA circuitry interacts with epileptic cortical neuron ensembles/engrams to generate seizure activity in the het *Gabrg2^Q390X^*mice. We have shown that the short activation of cortical epileptic neurons alone(supplemental Fig. 1, S1-ChR2 30 ms, simulating sleep spindle duration) cannot effectively trigger epileptic activity in the het *Gabrg2^Q390X^*KI mice. Only when NREM sleep state is induced by subcortical POA-ChR2 activation, the short activation of cortical epileptic neurons (S1-ChR2, 30ms) can effectively trigger epileptic activity with longer duration in the het *Gabrg2^Q390X^* KI mice (Fig. 5G/H, POA-ChR2/S1-ChR2). In contrast, when the POA activity is suppressed, the short activation of cortical epileptic neurons(S1-ChR2, 30 ms) becomes less effectively to trigger epilepsy and remaining epilepsy activity shows much shorter in the het *Gabrg2^Q390X^*KI mice (Fig. 6G/H, POA-NpHR/S1-ChR2). Moreover, this operating mechanism also contributes to myoclonic jerk generation in het *Gabrg2^Q390X^*KI mice, suggesting that sleep spindles^23,24,27,61^ within thalamocortical circuitry interact with other subcortical circuitry than thalamus to evolve into epileptic (poly)spike-wave discharges in epileptic patients. Consistent with this, MnPO/VLPO have been indicated to induce absence seizures in one WAG/Rjj rodent model^42^. In addition, we think that up-state dependent synaptic potentiation in cortical neurons

during NREM sleep contributes to underlying cellular mechanisms^41^ for this operational mechanism leading to sleep-related epilepsy incidence in animal models and human patients. As the subcortical POA functions as the sleep switch^2^, this operational mechanism can underlie sleep-related epilepsy incidence in refractory epilepsy^14,62,63^. Beyond this, this mechanism suggests the translational implication in epilepsy field. In treating genetic epilepsy and refractory epilepsy, we may have to suppress the up-state dependent synaptic potentiation within cortical neurons in epileptic patients to decrease the sleep-related seizure incidence, in addition to direct antiepileptic medicine to suppress cortical neuron excitability only^64–67^. This is indicated by the results of the POA suppression in the het *Gabrg2^Q390X^* KI mice (Fig. 6D-H for POA-NpHR/S1-ChR2), which essentially prevents sleep-related epilepitogenesis in the het *Gabrg2^Q390X^*KI mice [also myoclonic jerks(Fig. 7)]. One of drug 4-(diethylamino)-benzaldehyde (DEAB) to suppress up-state dependent synaptic potentiation in cortical neurons^41^ from the het *Gabrg2^Q390X^* KI mice can be used for this treatment. Compatible with this, DEAB administration (*i.p.*) does attenuate epilepsy incidence^21^ in our mouse het *Gabrg2^Q390X^*KI model (in its Fig.5 E-H^21^).

In addition, works on myoclonic seizures in mouse model have indicated that both somatosensory and motor cortex are involved to generate myoclonic jerks^47,48^. Although our cortex activation by blue laser focuses on the somatosensory cortex, due to reciprocally connection between the somatosensory and the motor cortex in the mouse brain^68–70^, epileptic activity within somatosensory cortex can influence motor cortex for myoclonic jerk generation^48^. Thus, this sleep-related seizure operational mechanism can contribute to myoclonic jerk generation which is implied by our data (Fig. 7B). However, we may need more experiments to support the subcortical POA circuitry involvement of myoclonic jerk generation within the motor cortex.

The sleep-dependent seizure incidence mechanism also influences the sleep spindle generation in epileptic patients. NREM sleep period is when sleep spindles occur and sleep-spindle dependent memories are consolidated in the brain^5,6,8,71^. And our data indicate that the POA nucleus is active within het *Gabrg2^Q390X^* KI mice (Fig. 1). In the wt mice, the POA nucleus activation by ChR2 can increase NREM sleep period (Fig. 4A/C) to promote sleep spindle generation in wt mice (supplemental Fig. 1) (also in our recent work^21^), while in the het *Gabrg2^Q390X^*KI mice, the POA nucleus activation by ChR2 increases NREM sleep period to trigger epileptic activity (Fig. 5E-H POA-ChR2/S1-ChR2). Most importantly, increased NREM sleep in the het *Gabrg2^Q390X^*KI mice does not accompany sleep spindle increase (supplemental Fig. 1) (also in our recent work, Fig. 6C^21^), unlike the sleep spindle increase in the wt mice^24,72^. It clearly suggests that, in the het *Gabrg2^Q390X^*KI mice during sleep period, epileptic cortical neurons seem to compete with nonepileptic/normal cortical neurons in the brain to generate ictal activity and suppress sleep spindle generation for memory consolidation^5,72^. Thus, in epileptic patients, suppressing sleep spindles’ for memory consolidation in the brain may explain cognitive deficits due to genetic and refractory epilepsy.

All together in this study, we show that the subcortical POA (VLPO and MnPO) sleep circuitry is active in the het *Gabrg2^Q390X^* KI mice and can effectively trigger epileptic activity in the het *Gabrg2^Q390X^* KI mice. This operational mechanism through the POA nucleus activation and epileptic cortical neuron activation underlies the sleep-related seizure incidences in genetic epilepsy and refractory epilepsy. Moreover, it also indicates the necessity to address sleep disturbance in refractory epilepsy patients for seizure treatment and this mechanism may contribute to memory/cognitive deficits in epileptic patients.

## ACKNOWLEDGMENTS

Chengwen Zhou designed experiments; Cobie Victoria Potesta, Madeleine Sandra Cargile and Chengwen Zhou performed experiments; Cobie Victoria Potesta provided animal care and surgery. Cobie Victoria Potesta, Madeleine Sandra Cargile and Chengwen Zhou analyzed data; Andrea Yan and Sarah Xiong did some imaging of brain sections. Chengwen Zhou wrote the manuscript and discussed with Drs. Martin J. Gallagher and Robert L. Macdonald.

This work was supported by NIH Grants NINDS R01NS107424-01 (Zhou) and NINDS/NIA R01 supplemental grant (Zhou).

All authors report no competing interests in relation to the work described.

## Figure Legend

**Supplemental Figure 1.** Short optogenetic activation of somatosensory cortex alone does not effectively trigger epileptic activity *in vivo* in het *Gabrg2^Q390X^* KI mice. Panels A/B show summary data of epileptic SWD/PSDs and their duration changes following somatosensory cortex ChR2 (S1-ChR2) activation. The inset in panel A shows the one representative experimental design for laser delivery. Wt n=7 mice, het n=7 mice. No significant changes with paired t-test.

## References

1. Saper, C.B., Chou, T.C. & Scammell, T.E. The sleep switch: hypothalamic control of sleep and wakefulness. Trends Neurosci 24, 726–731 (2001).

2. Saper, C.B., Fuller, P.M., Pedersen, N.P., Lu, J. & Scammell, T.E. Sleep state switching. Neuron 68, 1023–1042 (2010).

3. Saper, C.B., Scammell, T.E. & Lu, J. Hypothalamic regulation of sleep and circadian rhythms. Nature 437, 1257–1263 (2005).

4. Scammell, T.E., Arrigoni, E. & Lipton, J.O. Neural Circuitry of Wakefulness and Sleep. Neuron 93, 747–765 (2017).

5. Girardeau, G. & Lopes-Dos-Santos, V. Brain neural patterns and the memory function of sleep. Science 374, 560–564 (2021).

6. Ngo, H.V., Fell, J. & Staresina, B. Sleep spindles mediate hippocampal-neocortical coupling during long-duration ripples. Elife 9(2020).

7. Niethard, N., Ngo, H.V., Ehrlich, I. & Born, J. Cortical circuit activity underlying sleep slow oscillations and spindles. Proc Natl Acad Sci U S A 115, E9220–E9229 (2018).

8. Skelin, I., et al. Coupling between slow waves and sharp-wave ripples engages distributed neural activity during sleep in humans. Proc Natl Acad Sci U S A 118(2021).

9. Ahmed, O.J. & Vijayan, S. The roles of sleep-wake states and brain rhythms in epileptic seizure onset. J Neurosci 34, 7395–7397 (2014).

10. Beenhakker, M.P. & Huguenard, J.R. Neurons that fire together also conspire together: is normal sleep circuitry hijacked to generate epilepsy? Neuron 62, 612–632 (2009).

11. Bergmann, M., et al. A prospective controlled study about sleep disorders in drug resistant epilepsy. Sleep Med 75, 434–440 (2020).

12. Buchanan, G.F., et al. Proceedings of the Sleep and Epilepsy Workshop: Section 3 Mortality: Sleep, Night, and SUDEP. Epilepsy Curr 21, 15357597211004556 (2021).

13. Chauvette, S., Soltani, S., Seigneur, J. & Timofeev, I. In vivo models of cortical acquired epilepsy. J Neurosci Methods 260, 185–201 (2016).

14. Dell, K.L., et al. Seizure likelihood varies with day-to-day variations in sleep duration in patients with refractory focal epilepsy: A longitudinal electroencephalography investigation. EClinicalMedicine 37, 100934 (2021).

15. Gibbon, F.M., Maccormac, E. & Gringras, P. Sleep and epilepsy: unfortunate bedfellows. Arch Dis Child 104, 189–192 (2019).

16. Liguori, C., et al. Sleep disorders and late-onset epilepsy of unknown origin: Understanding new trajectories to brain amyloidopathy. Mech Ageing Dev 194, 111434 (2021).

17. Manokaran, R.K., Tripathi, M., Chakrabarty, B., Pandey, R.M. & Gulati, S. Sleep Abnormalities and Polysomnographic Profile in Children with Drug-Resistant Epilepsy. Seizure 82, 59–64 (2020).

18. Pavlova, M.K., et al. Proceedings of the Sleep and Epilepsy Workgroup: Section 2 Comorbidities: Sleep Related Comorbidities of Epilepsy. Epilepsy Curr, 15357597211004549 (2021).

19. Bagshaw, A.P., Rollings, D.T., Khalsa, S. & Cavanna, A.E. Multimodal neuroimaging investigations of alterations to consciousness: the relationship between absence epilepsy and sleep. Epilepsy Behav 30, 33–37 (2014).

20. Caraballo, R.H., Pasteris, M.C., Portuondo, E. & Fortini, P.S. Panayiotopoulos syndrome and diffuse paroxysms as the first EEG manifestation at clinical onset: a study of nine patients. Epileptic Disord 17, 143–149 (2015).

21. Catron, M.A., et al. Sleep slow-wave oscillations trigger seizures in a genetic epilepsy model of Dravet syndrome. Brain Commun 5, fcac332 (2023).

22. Papale, L.A., et al. Altered sleep regulation in a mouse model of SCN1A-derived genetic epilepsy with febrile seizures plus (GEFS+). Epilepsia 54, 625–634 (2013).

23. Contreras, D. & Steriade, M. Cellular basis of EEG slow rhythms: a study of dynamic corticothalamic relationships. J Neurosci 15, 604–622 (1995).

24. Kandel, A. & Buzsaki, G. Cellular-synaptic generation of sleep spindles, spike-and-wave discharges, and evoked thalamocortical responses in the neocortex of the rat. J Neurosci 17, 6783–6797 (1997).

25. Timofeev, I. & Steriade, M. Cellular mechanisms underlying intrathalamic augmenting responses of reticular and relay neurons. J Neurophysiol 79, 2716–2729 (1998).

26. Steriade, M. & Contreras, D. Relations between cortical and thalamic cellular events during transition from sleep patterns to paroxysmal activity. J Neurosci 15, 623–642 (1995).

27. Huguenard, J.R. & McCormick, D.A. Thalamic synchrony and dynamic regulation of global forebrain oscillations. Trends Neurosci 30, 350–356 (2007).

28. Halasz, P., Terzano, M.G. & Parrino, L. Spike-wave discharge and the microstructure of sleep-wake continuum in idiopathic generalised epilepsy. Neurophysiologie Clinique-Clinical Neurophysiology 32, 38–53 (2002).

29. McCormick, D.A. & Contreras, D. On the cellular and network bases of epileptic seizures. Annu Rev Physiol 63, 815–846 (2001).

30. Gvilia, I., et al. The role of adenosine in the maturation of sleep homeostasis in rats. J Neurophysiol 117, 327–335 (2017).

31. Lim, A.S., et al. Sleep is related to neuron numbers in the ventrolateral preoptic/intermediate nucleus in older adults with and without Alzheimer’s disease. Brain 137, 2847–2861 (2014).

32. Lu, J., Greco, M.A., Shiromani, P. & Saper, C.B. Effect of lesions of the ventrolateral preoptic nucleus on NREM and REM sleep. J Neurosci 20, 3830–3842 (2000).

33. Battaglia, F.P., Sutherland, G.R. & McNaughton, B.L. Hippocampal sharp wave bursts coincide with neocortical “up-state” transitions. Learn Mem 11, 697–704 (2004).

34. Besing, G.K., St John, E.K., Potesta, C.V., Gallagher, M.J. & Zhou, C. Artificial sleep-like up/down-states induce synaptic plasticity in cortical neurons from mouse brain slices. Front Cell Neurosci 16, 948327 (2022).

35. Cary, B.A. & Turrigiano, G.G. Stability of neocortical synapses across sleep and wake states during the critical period in rats. Elife 10(2021).

36. Dehghani, N., et al. Dynamic Balance of Excitation and Inhibition in Human and Monkey Neocortex. Sci Rep 6, 23176 (2016).

37. Levenstein, D., Watson, B.O., Rinzel, J. & Buzsaki, G. Sleep regulation of the distribution of cortical firing rates. Curr Opin Neurobiol 44, 34–42 (2017).

38. Torao-Angosto, M., Manasanch, A., Mattia, M. & Sanchez-Vives, M.V. Up and Down States During Slow Oscillations in Slow-Wave Sleep and Different Levels of Anesthesia. Front Syst Neurosci 15, 609645 (2021).

39. Tukker, J.J., Beed, P., Schmitz, D., Larkum, M.E. & Sachdev, R.N.S. Up and Down States and Memory Consolidation Across Somatosensory, Entorhinal, and Hippocampal Cortices. Front Syst Neurosci 14, 22 (2020).

40. Shu, Y., Hasenstaub, A. & McCormick, D.A. Turning on and off recurrent balanced cortical activity. Nature 423, 288–293 (2003).

41. Zhang, C.Q., et al. Impaired State-Dependent Potentiation of GABAergic Synaptic Currents Triggers Seizures in a Genetic Generalized Epilepsy Model. Cereb Cortex 31, 768–784 (2021).

42. Suntsova, N., et al. A role for the preoptic sleep-promoting system in absence epilepsy. Neurobiol Dis 36, 126–141 (2009).

43. Reijmers, L.G., Perkins, B.L., Matsuo, N. & Mayford, M. Localization of a stable neural correlate of associative memory. Science 317, 1230–1233 (2007).

44. Chuhma, N., Tanaka, K.F., Hen, R. & Rayport, S. Functional connectome of the striatal medium spiny neuron. J Neurosci 31, 1183–1192 (2011).

45. Garcia-Cabrero, A.M., et al. Laforin and malin deletions in mice produce similar neurologic impairments. J Neuropathol Exp Neurol 71, 413–421 (2012).

46. Arain, F., Zhou, C., Ding, L., Zaidi, S. & Gallagher, M.J. The developmental evolution of the seizure phenotype and cortical inhibition in mouse models of juvenile myoclonic epilepsy. Neurobiol Dis 82, 164–175 (2015).

47. Ding, L. & Gallagher, M.J. Dynamics of sensorimotor cortex activation during absence and myoclonic seizures in a mouse model of juvenile myoclonic epilepsy. Epilepsia 57, 1568–1580 (2016).

48. Ding, L., Satish, S., Zhou, C. & Gallagher, M.J. Cortical activation in generalized seizures. Epilepsia (2019).

49. Yu, F.H., et al. Reduced sodium current in GABAergic interneurons in a mouse model of severe myoclonic epilepsy in infancy. Nat Neurosci 9, 1142–1149 (2006).

50. Levesque, M., Macey-Dare, A.D.B., Wang, S. & Avoli, M. Evolution of interictal spiking during the latent period in a mouse model of mesial temporal lobe epilepsy. Curr Res Neurobiol 2, 100008 (2021).

51. Levesque, M., Wang, S., Macey-Dare, A.D.B., Salami, P. & Avoli, M. Evolution of interictal activity in models of mesial temporal lobe epilepsy. Neurobiol Dis 180, 106065 (2023).

52. Mohns, E.J., Karlsson, K.A. & Blumberg, M.S. The preoptic hypothalamus and basal forebrain play opposing roles in the descending modulation of sleep and wakefulness in infant rats. Eur J Neurosci 23, 1301–1310 (2006).

53. Suntsova, N., Szymusiak, R., Alam, M.N., Guzman-Marin, R. & McGinty, D. Sleep-waking discharge patterns of median preoptic nucleus neurons in rats. J Physiol 543, 665–677 (2002).

54. Kang, J.Q., Shen, W., Zhou, C., Xu, D. & Macdonald, R.L. The human epilepsy mutation GABRG2(Q390X) causes chronic subunit accumulation and neurodegeneration. Nat Neurosci 18, 988–996 (2015).

55. Zhou, C., et al. Altered cortical GABAA receptor composition, physiology, and endocytosis in a mouse model of a human genetic absence epilepsy syndrome. J Biol Chem 288, 21458–21472 (2013).

56. Zhou, C., Lippman, J.J., Sun, H. & Jensen, F.E. Hypoxia-induced neonatal seizures diminish silent synapses and long-term potentiation in hippocampal CA1 neurons. J Neurosci 31, 18211–18222 (2012).

57. Barth, A.L., Gerkin, R.C. & Dean, K.L. Alteration of neuronal firing properties after in vivo experience in a FosGFP transgenic mouse. J Neurosci 24, 6466–6475 (2004).

58. Roy, D.S., et al. Brain-wide mapping reveals that engrams for a single memory are distributed across multiple brain regions. Nat Commun 13, 1799 (2022).

59. Burnsed, J., Skwarzynska, D., Wagley, P.K., Isbell, L. & Kapur, J. Neuronal Circuit Activity during Neonatal Hypoxic-Ischemic Seizures in Mice. Ann Neurol 86, 927–938 (2019).

60. Alam, M.A., Kumar, S., McGinty, D., Alam, M.N. & Szymusiak, R. Neuronal activity in the preoptic hypothalamus during sleep deprivation and recovery sleep. J Neurophysiol 111, 287–299 (2014).

61. Contreras, D., Destexhe, A. & Steriade, M. Spindle oscillations during cortical spreading depression in naturally sleeping cats. Neuroscience 77, 933–936 (1997).

62. Rashed, H.R., et al. Refractory epilepsy and obstructive sleep apnea: is there an association? Egypt J Neurol Psych 55(2019).

63. Romero-Osorio, O., Gil-Tamayo, S., Narino, D. & Rosselli, D. Changes in sleep patterns after vagus nerve stimulation, deep brain stimulation or epilepsy surgery: Systematic review of the literature. Seizure-Eur J Epilep 56, 4–8 (2018).

64. Brandt, C., Gastens, A.M., Sun, M., Hausknecht, M. & Loscher, W. Treatment with valproate after status epilepticus: effect on neuronal damage, epileptogenesis, and behavioral alterations in rats. Neuropharmacology 51, 789–804 (2006).

65. Johannessen, C.U. & Johannessen, S.I. Valproate: past, present, and future. CNS Drug Rev 9, 199–216 (2003).

66. Sommer, B.R., Mitchell, E.L. & Wroolie, T.E. Topiramate: Effects on cognition in patients with epilepsy, migraine headache and obesity. Ther Adv Neurol Disord 6, 211–227 (2013).

67. White, H.S., Smith, M.D. & Wilcox, K.S. Mechanisms of action of antiepileptic drugs. Int Rev Neurobiol 81, 85–110 (2007).

68. Hooks, B.M. Sensorimotor Convergence in Circuitry of the Motor Cortex. Neuroscientist 23, 251–263 (2017).

69. Suter, B.A. & Shepherd, G.M. Reciprocal interareal connections to corticospinal neurons in mouse M1 and S2. J Neurosci 35, 2959–2974 (2015).

70. Petrof, I., Viaene, A.N. & Sherman, S.M. Properties of the primary somatosensory cortex projection to the primary motor cortex in the mouse. J Neurophysiol 113, 2400–2407 (2015).

71. Eckert, M.J., McNaughton, B.L. & Tatsuno, M. Neural ensemble reactivation in rapid eye movement and slow-wave sleep coordinate with muscle activity to promote rapid motor skill learning. Philos Trans R Soc Lond B Biol Sci 375, 20190655 (2020).

72. Andrillon, T., et al. Sleep spindles in humans: insights from intracranial EEG and unit recordings. J Neurosci 31, 17821–17834 (2011).

