## Supplementary figures and images for "Preoptic area controls sleep-related seizure onset in a genetic epilepsy mouse model"

### supplemental Fig. 1

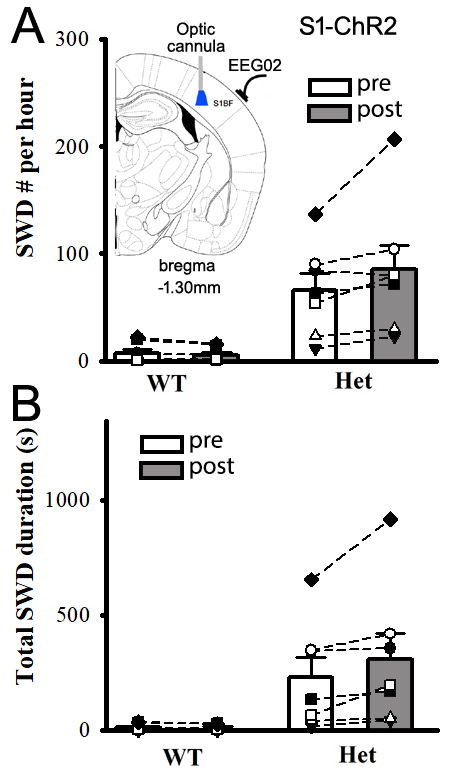
